# DDIT3 (CHOP) contributes to retinal ganglion cell somal loss but not axonal degeneration in DBA/2J mice

**DOI:** 10.1101/718882

**Authors:** Olivia J. Marola, Stephanie B. Syc-Mazurek, Richard T. Libby

## Abstract

Glaucoma is an age-related neurodegenerative disease characterized by the progressive loss of retinal ganglion cells (RGCs). Chronic ocular hypertension, an important risk factor for glaucoma, leads to RGC axonal injury at the optic nerve head. This insult triggers molecularly distinct cascades governing RGC somal apoptosis and axonal degeneration. The molecular mechanisms activated by ocular hypertensive insult that drive both RGC somal apoptosis and axonal degeneration are incompletely understood. The cellular response to endoplasmic reticulum stress and induction of pro-apoptotic DNA damage inducible transcript 3 (DDIT3, also known as CHOP) has been implicated as a driver of neurodegeneration in many disease models, including glaucoma. RGCs express DDIT3 upon glaucoma-relevant insults, and importantly, DDIT3 has been shown to contribute to both RGC somal apoptosis and axonal degeneration after acute induction of ocular hypertension. However, the role of DDIT3 in RGC somal and axonal degeneration has not been critically tested in a model of age-related chronic ocular hypertension. Here, we investigated the role of DDIT3 in glaucomatous RGC death using an age-related, naturally occurring ocular hypertensive mouse model of glaucoma, DBA/2J mice (D2). To accomplish this, a null allele of *Ddit3* was backcrossed onto the D2 background. Homozygous *Ddit3* deletion did not alter gross retinal or optic nerve head morphology, nor did it change the ocular hypertensive profile of D2 mice. In D2 mice, *Ddit3* deletion conferred mild protection to RGC somas but did not significantly prevent RGC axonal degeneration. Together, these data suggest that DDIT3 plays a minor role in perpetuating RGC somal apoptosis caused by chronic ocular hypertension-induced axonal injury, but does not significantly contribute to distal axonal degeneration.

## Introduction

Glaucoma is an age-related neurodegenerative disease characterized by the death of retinal ganglion cells (RGCs), the output neurons of the retina. An important risk factor for glaucomatous RGC death is elevated intraocular pressure (IOP), which leads to RGC axonal injury at the lamina cribrosa^1-5^ (termed the glial lamina in mice^5^). This insult is thought to trigger molecular signaling within RGCs that regulates somal degeneration proximal to the site of injury and axonal degeneration distal to the site of injury^6-10^. Identifying the molecular signaling pathways that lead from ocular hypertensive injury to RGC death is critical for understanding the pathobiology of glaucoma. To date, a mechanism important in both proximal and distal RGC degeneration has not been identified. The pro-apoptotic molecule BAX was shown to be required for RGC somal death but not axonal degeneration after chronic ocular hypertension and acute optic nerve injury^6,11,12^. Thus, the mechanism triggered by ocular hypertension that regulates glaucomatous neurodegeneration must ultimately converge upon BAX induction.

The adaptive response to endoplasmic reticulum (ER) stress (known as the unfolded protein response or the integrated stress response) has been implicated as a driver of neuronal death in neurodegenerative diseases, including glaucoma^13-17^. Upon prolonged and unresolved ER stress, the unfolded protein response has been shown to promote apoptosis via induction of DNA damage inducible transcript 3 (DDIT3, also known as CHOP). DDIT3 has been shown to act as a pro-apoptotic transcription factor; DDIT3 promoted transcription of pro-apoptotic *Bbc3*^18,19^, *Bim*^18,19^, *Gadd34*^20^, *Dr5*^19,21^, and *Ero1α*^22^ genes and inhibited transcription of the pro-survival gene *Bcl2*^19,20,23,24^. DDIT3 was also shown to be important for the translocation of activated BAX from the cytosol to the mitochondria^25,26^; allowing the intrinsic apoptotic cascade to ensue. Therefore, as a pro-apoptotic transcription factor upstream of BAX, DDIT3 may be an important regulator of RGC death upon glaucomatous insult.

DDIT3 has been shown to regulate RGC death in glaucoma and various other neurodegenerative diseases^27,28^. DDIT3 was expressed by RGCs after glaucoma-relevant insults, including optic nerve crush^13-15,29^ and the microbead model of acute ocular hypertension^13,14^. In addition, *Ddit3* was upregulated in both the retinas and optic nerve heads of mice with chronic ocular hypertension prior to the onset of glaucomatous neurodegeneration^30-32^. *Ddit3* deficiency or silencing was protective to RGC somas after mechanical axonal injury (optic nerve crush)^14,17,33^ and the microbead model of acutely induced ocular hypertension^14,33^. Interestingly, despite not appearing to have a major role in RGC axonal degeneration after optic nerve crush^17^, DDIT3 deficiency lessened axonal degeneration in an acute ocular hypertension model^33^. This protection, though minor, appeared roughly equal to the level of somal protection, suggesting that in some cells, *Ddit3* deficiency completely protected the RGC after an ocular hypertensive injury^33^.

DDIT3 appears to be an important mediator of RGC viability after glaucoma-relevant injuries. However, the role of DDIT3 in glaucomatous neurodegeneration has not been tested in a model of stochastic, age-related ocular hypertension. Here, we critically tested the role of DDIT3 in RGC axonal degeneration and somal loss in an inherited, age-related mouse model of chronic ocular hypertension. We found DDIT3 plays a minor role in RGC somal death but not axonal degeneration in the DBA/2J (D2) mouse model of chronic, age-related ocular hypertension^3,5,34-36^.

## Materials and Methods

### Mice

DBA/2J (D2) mice and mice with a null allele of *Ddit3*^37^ (B6.129S(Cg)-*Ddit3*^*tm2.1Dron*^/J) were obtained from the Jackson Laboratory (Stock numbers 000671 and 005530, respectively). The *Ddit3* null allele was backcrossed to the D2 background 10 times (>99% D2). After this backcross was completed, the D2.*Ddit3* colony was maintained by D2.*Ddit3*^*+/-*^ x D2.*Ddit3*^*+/-*^ intercrossing. D2.*Ddit3*^*+/+*^ environment-matched littermates were used as genetic controls for D2.*Ddit3*^-/-^ mice, and each genotype group included roughly equal numbers of females and males (D2.*Ddit3*^*+/+*^: 30 female, 34 male; D2.*Ddit3*^*-/-*^: 29 female, 31 male). Mice were fed chow and water *ad libitum* and were housed on a 12-hour light-to-dark cycle. All experiments were conducted in adherence with the Association for Research in Vision and Ophthalmology’s statement on the use of animals in ophthalmic and vision research and were approved by the University of Rochester’s University Committee on Animal Resources.

### Retina processing for plastic sectioning

As previously described^9,17,38,39^, eyes were enucleated and fixed for 24 hours in a solution of 2.5% glutaraldehyde, 2% paraformaldehyde (PFA) in 1X phosphate buffered saline (PBS; BioRad, 161-0780) at 4°C. Eyes were washed in 0.1M PO4, dehydrated in 50% ethanol for 1 hour, and placed in 70% ethanol overnight at 4 °C. Eyes were incrementally dehydrated in 80%, 95%, and 100% ethanol for one hour each at room temperature. Eyes were placed in acetone for 1 hour, washed with 100% ethanol for 1 hour, and placed in 1:1 100% ethanol: Hardener 1 Technovit 7100 (Electron Microscopy Sciences 14653) overnight at 4 °C. Eyes were then placed in Hadner I Technovit 7100 for 24 hours at 4 °C. Eyes were then incubated in 15:1 Hardener 1 Technovit 7100: Hardener 2 Technovit 7100 for 10 minutes on ice. Eyes were submerged in 15:1 Hardener 1 Technovit 7100: Hardener 2 Technovit 7100 and were allowed to harden in a plastic mold at room temperature. 2.5μm coronal cross sections were cut and collected on microscope slides. Sections that included the optic nerve head were stained with Multiple Stain Solution (Polysciences, Inc, 08824) for 1-2 minutes, washed with 100% ethanol, and cover-slipped with Permount (Fisher Scientific, SP15-500).

### Optic nerve processing for plastic sectioning and grading

Optic nerves were harvested and processed as previously described^6,9,38^. In brief, optic nerves were fixed *in situ* in 2.5% glutaraldehyde, 10% formalin in 1X PBS for 24 hours at 4 °C. Nerves were dissected from the brain and were incubated in 1% osmium for 2 hours at room temperature. Otherwise, nerves were processed identically to eyes as described above. 1.5*μ*m cross sections were cut and collected on microscope slides. Nerve sections were stained with 1% paraphenylenediamine (PPD) in absolute methanol for 15 minutes, and washed with 100% ethanol for 10 minutes. PPD stains the myelin sheath of all axons but differentially darkly stains the axoplasm of dying axons. A masked observer used a validated grading scale to assess the level of glaucomatous damage of each optic nerve. As previously described^5,6,9,40^, nerves with <5% axons damaged or lost (consistent with axonal loss associated with normal aging) were judged to have no/early damage, nerves judged to have moderate damage had 5-50% axonal damage or loss (averaging ∼30% loss) often with localized areas of gliosis, and nerves with >50% axonal damage or loss, often with large areas of glial scaring, were judged to have severe damage. A masked observer selected optic nerves with the most axonal damage (judged to have <5% axonal survival) for assessment of RGC somal survival.

### Controlled optic nerve crush

Controlled optic nerve crush (CONC) was performed as previously described^6,38,39^. Briefly, mice were anesthetized with intraperitoneal 100mg/kg ketamine and 10mg/kg xylazine. Analgesic 2mg/kg meloxicam was administered subcutaneously prior to surgery. The optic nerve was exposed and crushed immediately behind the eye with self-closing forceps for 5 seconds. Sham surgery was performed on the contralateral eye, where the optic nerve was exposed but not crushed. Antibiotic ointment was applied to the eyes following the procedure. Eyes were harvested 5 and 14 days post-CONC.

### Intraocular pressure measurement

As previously described^9,35,38^, intraocular pressures were measured by a masked observer using Tonolab (Colonial Medical Supply, Franconia, NH, USA) according to manufacturer’s instructions 3-5 minutes after intraperitoneal administration of anesthetic 100mg/kg ketamine and 10mg/kg xylazine.

### Immunofluorescence

As previously described^38,39^, eyes were harvested and fixed in 4% paraformaldehyde in 1X PBS for 90 minutes. Retinas were dissected free from the optic cup and blocked in 10% horse serum, 0.4% Triton™ X-100 (Fisher scientific, 9002-93-1) in 1X PBS overnight at 4 °C. Retinas were then incubated at 4 °C for 3 days in primary antibody diluted in 10% horse serum, 0.4% Triton™ X-100 in 1X PBS. Retinas were then washed and incubated for 24 hours at 4 °C in secondary antibodies diluted in 1X PBS. Retinas were washed and mounted on microscope slides ganglion cell layer-up in Flourogel in TRIS buffer (Electron Microscopy Sciences, 17985-11).

**Table 1.**
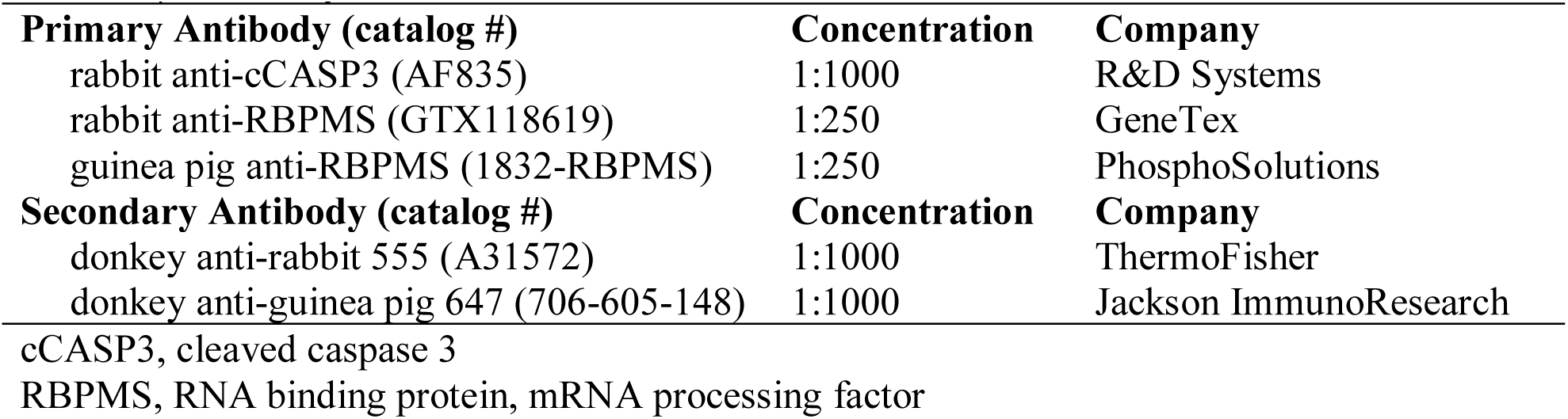
Summary of antibodies.

### Cell quantification

As previously described^38,39^, cCASP3+ RBPMS+ cells were quantified using eight 20x fields per retina, and RBPMS+ cell counts were assessed using eight 40x fields per retina. Images were taken approximately 220μm from the peripheral edge of the retina and were equally spaced from each other. The manual cell counter plug-in in ImageJ was utilized for cell quantification. Retinal imaging and cell quantifications were performed by a masked observer. Cell quantifications were normalized to the total area measured and reported as cells/mm^2^.

### Statistical analysis

Data were analyzed using GraphPad Prism8 software. Comparisons between two groups (cCASP3+ cells/mm^2^ after CONC between genotypes, Fig. 1a and %RGCs in retinas with severe optic nerves between genotypes, Fig. 4) were analyzed using an unpaired two-tailed student’s t test. Comparisons across more than two groups (RGCs/mm^2^ 14 days after sham and CONC procedures between genotypes, Fig. 1b) or two groups across multiple timepoints (IOP measurements at multiple timepoints between genotypes, Fig. 2b) were analyzed using a two-way ANOVA followed by a Sidak *post hoc* test. For these statistical tests, multiplicity adjusted P values are reported. The comparison of the percent of optic nerves at each grade between genotypes (Fig. 3b) was analyzed using a Chi-square test. P values of <0.05 were considered statistically significant. Throughout the manuscript, results are reported as mean± standard error of the mean (SEM).

**Fig. 1.**
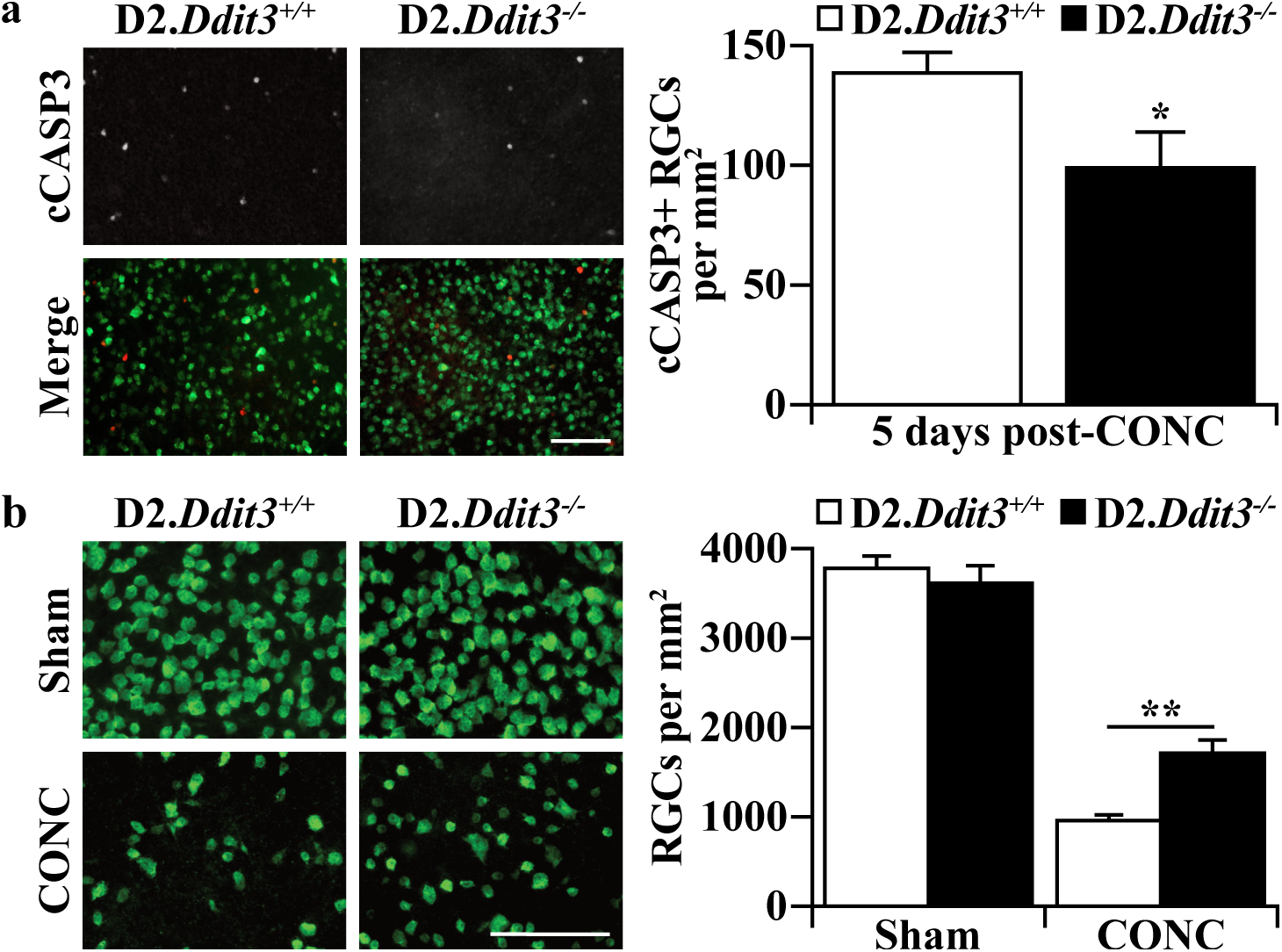
D2 background does not alter protection conferred by *Ddit3* deletion after axonal injury. **a)** Retinal flat mounts 5 days post-CONC stained for the RGC marker RBPMS (green) and cleaved caspase 3 (cCASP3, red). There are significantly fewer cCASP3+ RGCs (CASP3+RBPMS+ cells) in D2.*Ddit3*^*-/-*^ retinas compared with D2.*Ddit3*^*+/+*^ retinas (cells/mm^2^±SEM: D2.*Ddit3*^*+/+*^, 139.7±7.5, D2.*Ddit3*^*-/-*^, 100.2±13.8; n=6 per genotype, P=0.031, two-tailed t test). **b)** Retinal flat mounts 14 days post-CONC stained for RBPMS. D2.*Ddit3*^*+/+*^ and D2.*Ddit3*^*-/-*^ retinas had similar RGC densities 14 days post-sham surgery (cells/mm^2^±SEM: D2.*Ddit3*^*+/+*^, 3811.4±109.5, D2.*Ddit3*^*-/-*^, 3647.0±166.2; n=6 per genotype; P=0.910, two-way ANOVA, Sidak *post hoc*). Both genotypes had a significant reduction of RGCs 14 days post-CONC (P<0.001 for both comparisons, n=6 per genotype, two-way ANOVA, Sidak *post hoc*), however, D2.*Ddit3*^*-/-*^ retinas had significantly more surviving RGC compared to D2.*Ddit3*^*+/+*^ controls (cells/mm^2^±SEM: D2.*Ddit3*^*+/+*^, 991.0±33.9, D2.*Ddit3*^*-/-*^, 1745.8±116.5; n=6 per genotype; P=0.001, two-way ANOVA, Sidak *post hoc*). Error bars, SEM. Scale bars, 100μm.

**Fig. 2.**
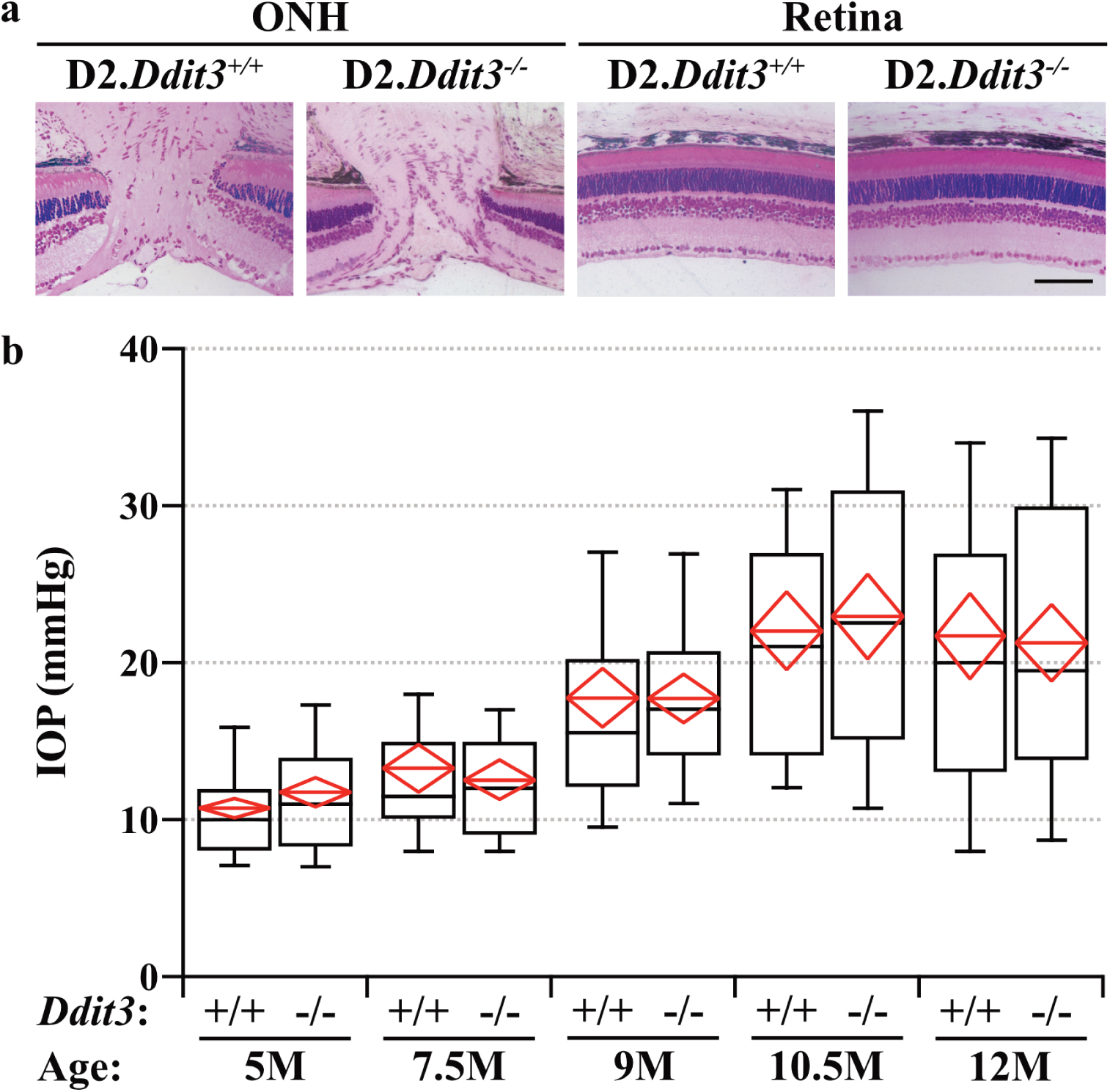
*Ddit3* deletion does not alter D2-associated endophenotypes. **a)** Semi thin plastic sections of D2.*Ddit3*^*+/+*^ and D2.*Ddit3*^*-/-*^ ONHs and retinas. *Ddit3* deficiency did not cause gross morphological abnormalities of the ONH (left) or retina (right) in D2 mice (n=6 per genotype). **b)** D2.*Ddit3*^*+/+*^ and D2.*Ddit3*^*-/-*^ IOPs taken at 5M (n≥76), 7.5M (n≥72), 9M (n≥60), 10.5M (n≥56), and 12M (n≥66). There was no difference in IOP between D2.*Ddit3*^*+/+*^ and D2.*Ddit3*^*-/-*^ eyes at any time point (P>0.05 for all comparisons, two-way ANOVA, Sidak *post hoc*). Note, for both genotypes, 9M, 10.5M, and 12M IOPs were significantly elevated compared to the 5M timepoint (P<0.001 for all comparisons, two-way ANOVA, Sidak *post hoc*). The black line represents the median, and the upper and lower bounds represent the 90^th^ and 10^th^ percentiles, respectively. The red line represents the mean, and the top and bottom points of the red diamond represent the 95% confidence interval. Scale bar, 100μm.

**Fig. 3.**
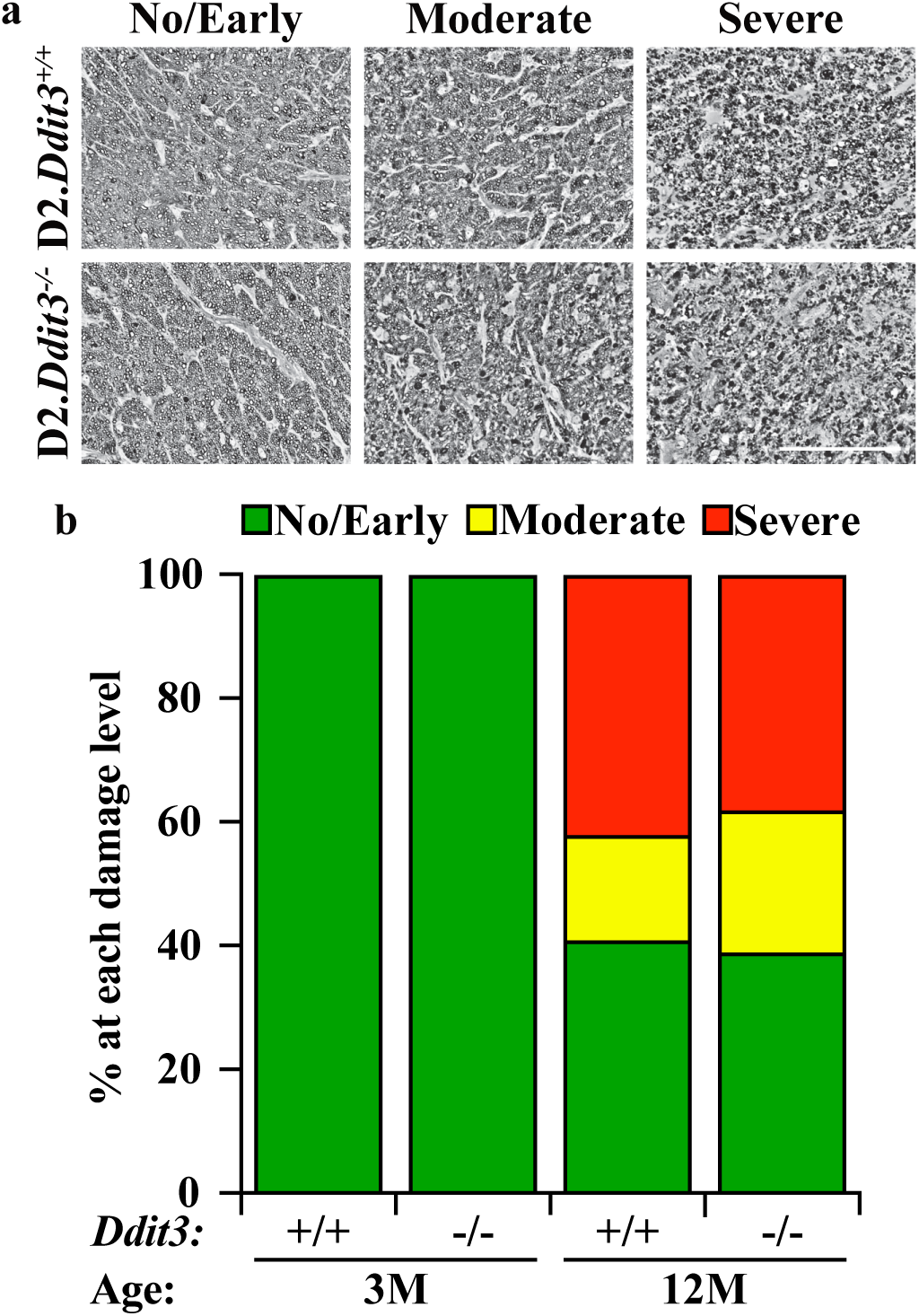
*Ddit3* deletion does not prevent RGC axon degeneration in a model of chronic ocular hypertension. **a)** Representative optic nerve cross sections stained with PPD from 12M D2.*Ddit3*^+/+^ and D2.*Ddit3*^-/-^ mice judged to have no/early (left), moderate (middle) and severe (right) axonal damage. **b)** Percentages of D2.*Ddit3*^*+/+*^ and D2.*Ddit3*^*-/-*^ optic nerves judged to have no/early, moderate, and severe axonal damage at 3M and 12M. D2.*Ddit3*^*+/+*^ and D2.*Ddit3*^*-/-*^ optic nerves had no moderate or severe damage at 3M (n=10 per genotype). D2.*Ddit3*^*+/+*^ and D2.*Ddit3*^*-/-*^optic nerves had similar levels of axonal damage at 12M (D2.*Ddit3*^*+/+*^, D2.*Ddit3*^*-/-*^: no/early, 41%, 40%; moderate, 17%, 23%; severe, 42%, 37%, n≥60 per genotype, P=0.474, Chi-square test). Scale bar, 100μm.

**Fig 4.**
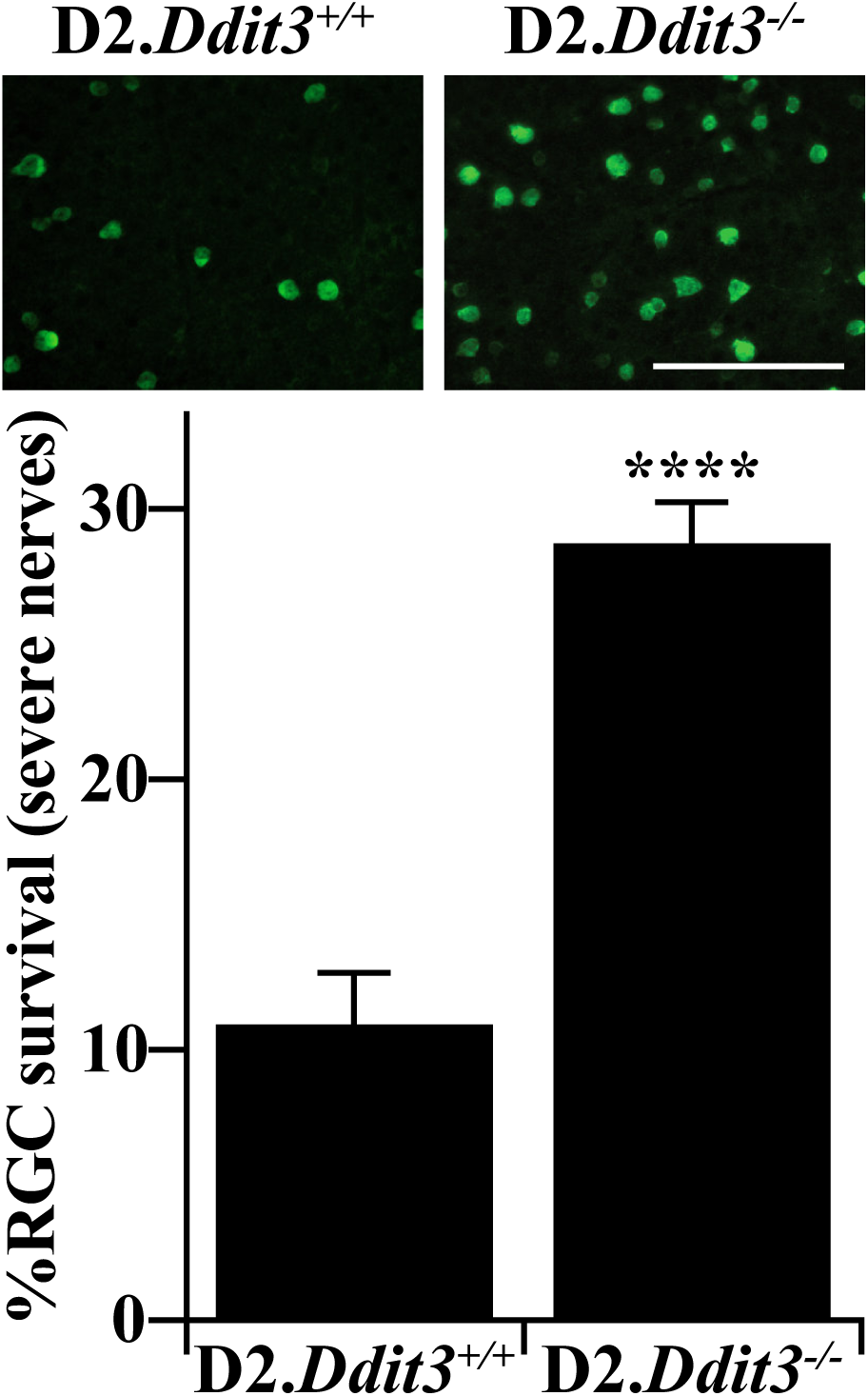
DDIT3 contributes to chronic ocular hypertension-induced RGC somal degeneration. RBPMS+ RGC somal density was assessed for 12M D2.*Ddit3*^*+/+*^ *and* D2.*Ddit3*^*-/-*^ retinas with corresponding optic nerves judged to have the most severe levels of axonal degeneration (judged to have < 5% axons remaining). D2.*Ddit3*^-/-^ retinas had significantly more surviving RGC somas compared to D2.*Ddit3*^*+/+*^ controls with equally severe optic nerves (%survival±SEM; D2.*Ddit3*^*+/+*^: 11.0±1.8%, D2.*Ddit3*^*-/-*^: 28.8±1.4%, n≥10, P<0.001, two-tailed t test). RGC counts were normalized to young (1.5-2.5M) control RGC counts of the respective genotype, see Fig. 1b. Error bars, SEM. Scale bar, 100μm.

## Results

### D2 background does not alter protection conferred by *Ddit3* deletion after axonal injury

*Ddit3* deficiency on the C57BL/6J background provided protection to some RGC somas after controlled optic nerve crush (CONC)^14,17,33^. However, genetic background may affect RGC death after axonal insult^41,42^. To determine if *Ddit3* deficiency lessened RGC death after axonal injury on the DBA/2J (D2) genetic background, CONC was performed on young (1.5-2.5 months of age, M) D2 mice. At this age, wildtype D2 mice do not yet have elevated IOP or any morphological glaucomatous damage^35^. D2.*Ddit3*^*-/-*^ and D2.*Ddit3*^*+/+*^ retinas were harvested 5 and 14 days after CONC. Consistent with CONC-induced RGC death in C57BL/6J mice, D2.*Ddit3*^*-/-*^ retinas had 28.1% fewer cleaved caspase 3+ (cCASP3; cleavage of CASP3 is a critical step of apoptosis) RGCs compared to D2.*Ddit3*^*+/+*^ retinas 5 days post-CONC (Fig. 1a). D2.*Ddit3*^*+/+*^ and D2.*Ddit3*^*-/-*^ retinas had similar RGC densities 14 days post-sham surgery (Fig. 1b) as judged by a specific marker for RGCs (RNA binding protein, mRNA processing factor; RBPMS)^43,44^. Retinas of both genotypes had significant RGC loss 14 days post-CONC as compared to sham, however, D2.*Ddit3*^*-/-*^retinas had 29.6% increased RGC survival compared to D2.*Ddit3*^*+/+*^ controls (Fig. 1b). Thus, on the D2 genetic background, *Ddit3* deficiency provided similar protection to RGCs as previous reports using the C57BL/6J genetic background after axonal injury^14,17,33^.

### *Ddit3* deletion does not alter D2-associated endophenotypes

The optic nerve head (ONH) is an important site in the pathobiology of glaucoma. In ocular hypertensive DBA/2J mice, the ONH is likely the site of an early critical axonal injury^4,5^. To ensure *Ddit3* deletion did not cause any developmental retinal or ONH abnormalities in D2 mice, D2.*Ddit3*^*-/-*^ and D2.*Ddit3*^*+/+*^ retinal and ONH morphology were assessed at 1.5-3M. *Ddit3* deletion caused no gross morphological retinal or optic nerve head abnormalities in D2 mice as judged by semi-thin sections (Fig. 2a).

ER stress has been implicated in regulating IOP elevation for some genetic causes of glaucoma^45,46^. Since RGC degeneration in D2 mice depends on age-related IOP elevation^36,47-51^, it was important to determine if *Ddit3* deficiency altered IOP elevation in D2 mice. IOP was assessed at 5M, 7.5M, 9M, 10.5M, and 12M (Fig. 2b). As a population, IOP was not elevated at 5M and 7.5M. Both genotypes had significant IOP elevations at 9M, 10.5M, and 12M compared to baseline IOPs taken at 5M. D2.*Ddit3*^*-/-*^ mice had similar IOPs to D2.*Ddit3*^*+/+*^ mice at each timepoint measured, thus, *Ddit3* deletion did not alter the stereotypic IOP profile of D2 mice.

### *Ddit3* deletion does not significantly prevent RGC axonal degeneration in a model of chronic ocular hypertension

DDIT3 has been implicated in driving both RGC somal and axonal degeneration after acute axonal injury^52^ and acute ocular hypertension^33^. To determine whether *Ddit3* regulated glaucomatous neurodegeneration in a chronic model of age-related ocular hypertension, D2.*Ddit3*^*-/-*^ and D2.*Ddit3*^*+/+*^ littermate control optic nerves were assessed for glaucomatous damage at 12M. At this time point, a significant proportion of D2 optic nerves have severe levels of axonal degeneration^9,35,38^. Optic nerves from young (1.5-3M) D2.*Ddit3*^*+/+*^ and D2.*Ddit3*^*-/-*^ mice were also assessed to ensure no premature glaucomatous damage or axonal phenotype occurred in D2.*Ddit3*^*-/-*^ mice. Optic nerve damage was graded as “no/early”, “moderate” or “severe” using a validated grading scale^5,6,9,40^ (Fig. 3a, see methods for grading details). Neither D2.*Ddit3*^*+/+*^ nor D2.*Ddit3*^*-/-*^ optic nerves exhibited any signs of axonal degeneration at 1.5-3M (Fig. 3b). At 12M, D2.*Ddit3*^*-/-*^ mice had similar levels of optic nerve damage compared to D2.*Ddit3*^*+/+*^ controls (Fig. 3b). Therefore, *Ddit3* deletion did not provide protection to RGC axons in D2 mice, suggesting that *Ddit3* is likely not a critical regulator of axonal degeneration in a model of chronic age-related ocular hypertension.

### DDIT3 contributes to chronic ocular hypertension-induced RGC somal degeneration

Because RGC distal axonal degeneration and proximal somal apoptosis are regulated by molecularly distinct pathways upon axonal insult^6,8-10,17,53^, it was important to determine whether DDIT3 governs proximal RGC somal apoptosis in a model of age-related ocular hypertension. To accomplish this, retinas judged to have the most severe levels of optic nerve degeneration (<5% axons remaining) from both genotypes were assessed for RGC somal survival. While D2.*Ddit3*^*+/+*^ retinas with corresponding severe optic nerves had only 11.0±1.8% somal survival, D2.*Ddit3*^*-/-*^ retinas had 28.8±1.4% somal survival (Fig. 4), consistent with levels of protection conferred by *Ddit3* deletion after mechanical axonal injury (Fig. 1b and^17^). Therefore, DDIT3 plays a minor role in chronic ocular hypertension-induced RGC somal apoptosis.

## Discussion

Chronic ocular hypertension is an important risk factor for the development of glaucomatous neurodegeneration. Ocular hypertension is thought to injure RGCs as they exit the eye at the lamina cribrosa^1-5^. Axonal injury is thought to trigger both RGC somal and axonal degeneration pathways^6-10^. ER stress, specifically DDIT3, has been implicated as a driver of RGC death after glaucoma-relevant injuries^13-15,17,33^. Importantly, DDIT3 was shown to regulate both RGC axonal degeneration and somal apoptosis in models of mechanical axonal injury and acutely-induced ocular hypertension^33^. In the present work, the role of DDIT3 in age-related, chronic ocular hypertension-induced RGC death was investigated. While *Ddit3* deletion in D2 mice provided mild protection to RGC somas, it did not significantly prevent RGC axonal degeneration. These data suggested DDIT3 has a minor role in regulating RGC somal death after axonal injury induced by chronic ocular hypertension. Therefore, the molecular process triggered by ocular hypertension that governs both RGC somal apoptosis and axonal degeneration remains unknown.

DDIT3 played a minor role in RGC somal degeneration in ocular hypertensive D2 mice; *Ddit3* deletion protected ∼20% of RGC somas in retinas with severe optic nerve axonal degeneration. The pro-apoptotic molecule BAX was shown to be required for RGC somal degeneration after CONC^6,54^ and in ocular hypertensive D2 mice^6^, however, BAX does not regulate RGC axonal degeneration in these models^6^. DDIT3 is important in the translocation of BAX from the cytosol to the mitochondria during prolonged ER stress^25,26^. However, because *Ddit3* deletion only protects ∼20% of RGC somas in D2 mice, another mechanism must work in tandem with DDIT3 to induce BAX. The mitogen-activated protein kinase effector and transcription factor JUN was shown to be an important regulator of ocular hypertension-induced RGC somal apoptosis. In fact, *Jun* deficiency protected ∼2.5 times more RGC somas compared to *Ddit3* deficiency in 12M D2 mice with severe optic nerve degeneration^9^. Interestingly, JUN and DDIT3 additively contribute to RGC somal apoptosis after CONC; dual deletion of *Jun* and *Ddit3* conferred 75% somal protection 120 days post-CONC (*Jun* and *Ddit3* deletion alone allowed 48% and 25% protection at this timepoint, respectively)^17^. Therefore, identifying the ocular hypertension-induced upstream regulator of both JUN and DDIT3 may be an important step in determining an upstream mechanism driving glaucomatous RGC death. Further, the role of both *Jun* and *Ddit3* in glaucomatous neurodegeneration should be tested in a model of age-related chronic ocular hypertension.

Previous reports have shown that DDIT3 deficiency lessened both RGC axonal degeneration and somal loss in the microbead model of acute IOP elevation and after CONC^33^. However, we report no gross difference in RGC axonal degeneration between D2.*Ddit3*^*+/+*^ and D2.*Ddit3*^*-/-*^ mice at 12M. This result is consistent with our previous report that *Ddit3* deletion does not protect from loss of RGC axonal conductance after CONC in C57BL/6J mice^17^, unlike manipulation of molecules known to protect axons from degeneration (*Wld*^*S*^ and *Sarm1*^10^). The differences between these results could perhaps be explained by the nature and/or duration of the insults. In the microbead model of acute ocular hypertension, optic nerves had only moderate neurodegeneration (∼29% axonal loss) after 8 weeks^33^. It is possible that DDIT3 deficiency can delay axonal degeneration after an ocular hypertensive injury, but not prevent degeneration after long term ocular hypertensive insult or severe mechanical injury. It is also conceivable that the differences in findings are explained by the age-related nature of the DBA/2J disease, as acute induction of ocular hypertension was performed on young animals^33^. Finally it is possible that there is a small number of axons surviving in the D2.*Ddit3*^*-/-*^ optic nerve that were not detected using a grading system. Regardless, our findings suggest that DDIT3 does not play a major role in axonal degeneration in an age-related, chronic ocular hypertension model of glaucoma.

In conclusion, the role of DDIT3 in glaucomatous neurodegeneration was tested in an inherited model of chronic, age-related ocular hypertension. DDIT3 deficiency did not alter retinal or optic nerve morphology, nor did it alter the IOP profile of the D2 model. In this model, DDIT3 does not contribute to RGC axonal degeneration signaling, but it is responsible for ∼20% of RGC somal apoptosis. Future work should focus on the roles of both JUN and DDIT3 together in perpetuating glaucomatous RGC death, and should elucidate upstream regulators of both JUN and DDIT3 after glaucoma-relevant injury.

## Acknowledgements

The authors would like to acknowledge Alyssa West and Thurma McDaniel for their excellent technical support. This work was supported by EY018606 (RTL), Research to Prevent Blindness, an unrestricted grant to the Department of Ophthalmology at the University of Rochester Medical Center, the NIH Institutional MSTP Training Grant T32 GM007356 (SBSM), and the NEI of the NIH under Award Number T32, EY007125 (OJM). The content is solely the responsibility of the authors and does not necessarily represent the official views of the NIH. The funding agencies had no role in the design of the study and collection, analysis, and interpretation of data and in writing the manuscript.

